# Mating strategy is determinant of Adenovirus prevalence in European bats

**DOI:** 10.1101/626309

**Authors:** Federica Rossetto, Maria Iglesias-Caballero, H. Christoph Liedtke, Ivan Gomez-Mestre, Jose M. Berciano, Gonzalo Pérez-Suárez, Oscar de Paz, Juan E. Echevarría, Inmaculada Casas, Javier Juste

## Abstract

Adenoviruses are double-strained DNA viruses found in a great number of vertebrates, including humans. In order to understand their transmission dynamics, it is crucial, even from a human health perspective, to investigate how host traits influence their prevalence. Bats are important reservoirs for Adenoviruses, and here we use the results of recent screenings in Western Europe to evaluate the association between characteristic traits of bat species and their probability of hosting Adenoviruses, taking into account their phylogenetic relationships. Across species, we found an important phylogenetic component in the presence of Adenoviruses and mating strategy as the most determinant factor conditioning the prevalence of Adenoviruses across bat species. Contrary to other more stable mating strategies (e.g. harems), swarming could hinder transmission of Adenoviruses since this strategy implies that contacts between individuals are too short. Alternatively, bat species with more promiscuous behavior may develop a stronger immune system. Outstandingly high prevalence of Adenoviruses was reported for the Iberian species *Pipistrellus pygmaeus, P. kuhlii* and *Nyctalus lasiopterus* and we found that in the latter, males were more likely to be infected by Adenoviruses than females, due to the immunosuppressing consequence of testosterone during the mating season. As a general trend across species, we found that the number of Adenoviruses positive individuals was different across localities and that the difference in prevalence between populations was correlated with their geographic distances (*P. pygmaeus*). These results increase our knowledge about the transmission mechanisms of Adenoviruses.

**Author Summary:** Adenoviruses are DNA viruses with a wide range of vertebrate hosts, including humans, causing ocular, respiratory and gastrointestinal diseases. Here, we focus on the prevalence of Adenoviruses in bats, which are known to be natural reservoir of many viruses, using the results of recent screenings for prevalence of these viruses in 33 European bat species. Our aim is to find association between Adenoviruses prevalence and biological and behavioral host traits, considering the heterogeneity both between and within species in order to have a deeper understanding of mechanisms of viral transmission.

Our results highlight the importance of mating strategy: bats species using swarming as mating strategy are less likely to be infected by Adenoviruses. Moreover, we found that locality of capture can explain a higher prevalence of Adenovirus within species. However, no general pattern has been found in the analysis at individual level, suggesting a strong species specificity and complex viral transmission dynamics.

## Introduction

Some viruses have the potential for cross-species transmission, with spillover episodes from humans to wildlife and vice versa, a phenomenon referred to as zoonosis [1]. Current research on the dynamics behind host/virus interactions and inter- and intra-specific transmissions is of scientific interest and important implications for emerging zoonoses and consequently for public health.

The heterogeneity (understood as the inter- and intra-specific variation) in the ability of hosts to transmit pathogens is among the most fundamental concepts in disease dynamics and crucial in the design of disease control strategies [2]. The potentiality of cross-species or cross-populations transmission depends on the characteristics of host and virus traits and is affected directly or indirectly by a variety of ecological, social, or socioeconomic changes [3] that can lead to new conditions boosting the expansion of the viruses to new host species or populations. Recent studies have revealed that traits such as host phylogenetic relatedness, conservation status, and geographical overlap are critical in the potential for cross-species transmission of viruses [4,5].

The highest proportion of zoonotic viruses in mammals is found in bats, primates and rodents [6]. Specific bat characteristics like their ability to fly, unique among mammals, or their migratory capacity [7], increase their potential role as vectors of diseases. Bats are natural reservoirs for many viruses, such as Coronavirus, Nipah virus and Ebola related to new emerging diseases that have received attention in the last years due to the potential risk they pose for human pandemic events [8–10] or with important health and economic consequences world-wide, such as rabies outbreaks [11,12]. The capacity to coexist with viruses (in the absence of disease) is probably linked to the bats’ immune-system which seems be different to that of other mammals, [7,13] and related to the development of altered mitochondrial genomes [14]. This pathogen control in bat hosts has favored ancient events of coevolution or parallel evolution [15,16] between bats and viruses.

Additionally, bats are systematically very diverse and form the second largest order of mammals [17]. In fact, bats occupy all kinds of habitats throughout all continents except Antarctica, showing a surprising ecological breadth that implies large variation in trait characteristics either in their morphology: e.g. size variation across species [18]; social behavior: e.g. differences in mating system [19]; or habitat requirements: e.g. roosts preferences from trees to caves [18]. For all these reasons, bats provide a good model to inspect the correlation of host trait characteristics with the prevalence of viruses.

Adenoviruses (AdVs) are non-enveloped dsDNA viruses with a broad range of vertebrate hosts, including humans. These viruses have been grouped into five genera [20]: *Mastadenovirus* (mammals), *Atadenovirus* (mammals, birds and reptiles), *Aviadenovirus* (birds), *Siadenovirus* (poultry and amphibians) and *Ichtadenovirus* (fish). Symptoms in humans include respiratory, eye infections and intestinal or digestive illness and seldom lead to mortality [21,22]. Since the first detection of an AdV in a fruit bat from Japan [23], a high diversity of AdVs has been isolated from bats from America, Africa and Asia, all grouped within the genus *Mastadenovirus* [24]. The first AdV in a European bat was isolated from a common pipistrelle (*Pipistrellus pipistrellus*) in Germany [25]. Later, Vidovszky *et al.* [26] obtained partial sequences from as many as 28 presumably new AdVs hosted by 12 different bat species in Germany and Hungary. Iglesias-Caballero *et al.* [16] conducted a country-wide survey in Spain constituting the largest screening for AdVs in bats to date, checking >1,000 individuals belonging to 28 species and focusing not only on the analysis of fecal samples and internal tissues -as in previous studies-but also on the analysis of the oropharyngeal swabs. AdVs are detected in almost half of the Iberian bat species studied, in both feces and in the upper respiratory tract (for the first time in bats), establishing a possible fecal-oral transmission route in two *Pipistrellus pygmaeus*. Moreover, they found a surprisingly high prevalence in this species together with the co-generic *P. kuhlii* and the closely related giant noctule, *Nyctalus lasiopterus*, which accounted for the vast majority of viruses detected in the study. These results have presented us with the opportunity to analyze the abiotic factors or biotic traits determining prevalence of AdVs in European bats at two levels: across-species and among individuals within a species.

The analysis of morphological and behavioral traits and presence of AdVs in bats is also useful for the understanding of viral transmission dynamics. In fact, little is known about the transmission mechanisms of AdVs and no study has focused on transmission in bats. A handful of studies in humans suggest that transmission may need direct contact or at least a droplet spraying (such as those produced by coughing or sneezing) or aerosol [27–29]. The strong host specificity and the parallelism between host and AdVs phylogenies found for bats [16,30] and for primates [31] strongly suggest that cross-species switching of mastadenovirus are not frequent events although they have been detected in AdVs evolution [30].

No study has focused so far on the analyses of the determinant factors influencing prevalence of viruses on bats at both species and individual levels. Across species, Webber *et al.* [32] found that viral richness was positively correlated with group size as predicted by the contact-rate hypothesis. In turn, at the intra-specific level Dietrich *et al.* [33] found an important seasonal shift in prevalence with a significant increase in AdVs shedding during reproduction while studying AdVs prevalence in two bats (*Miniopterus natalensis* and *Rousettus aegyptiacus*). The aim of this study is to advance our understanding of heterogeneity in the prevalence of AdVs in bats, testing the association between traits and presence of AdVs at both the among-species and within-species levels. Among species, and according to the contact-rate hypothesis, higher prevalence of AdVs is expected in species roosting in large groups, sharing refuge with other species and/or mating in swarming aggregations. In addition, we hypothesize a strong phylogenetic signal given the strong species-specificity found in the presence of AdVs. Within species, we predict a higher prevalence in females because of their concentration in large numbers in nursery colonies [34], with high contact rate and a high concomitant risk of infection.

In summary, our aims were to: 1) Test for significant phylogenetic component to the presence of AdVs in European bats; 2) test the importance of ecological characteristics at a species-level for the presence of AdVs taking into account phylogenetic relationships; 3) investigate whether some individual characteristics are determinant to explain the differences in the prevalence of AdVs within species.

## Results

The final working database consisted of 1,985 bats sampled and checked for AdVs, belonging to 10 genera (*Barbastella, Eptesicus, Hypsugo, Miniopterus, Myotis, Nyctalus, Pipistrellus, Plecotus, Rhinolophus, Vespertilio*) and representing 33 of the 45 European bat species. This database included a total of 1,612 bats belonging to 27 Iberian species surveyed for the study (Fig 1). The inclusion in the analyses of the published results from Vidovszky *et al.* [26] and Sonntag *et al.* [25] for Germany and Hungary, allowed the addition of seven European species (*Eptesicus nilssonii, Myotis brandtii, M. dasycneme, M. nattereri, Pipistrellus nathusii* and *Vespertilio murinus*), not included in the Iberian database.

**Fig 1.**
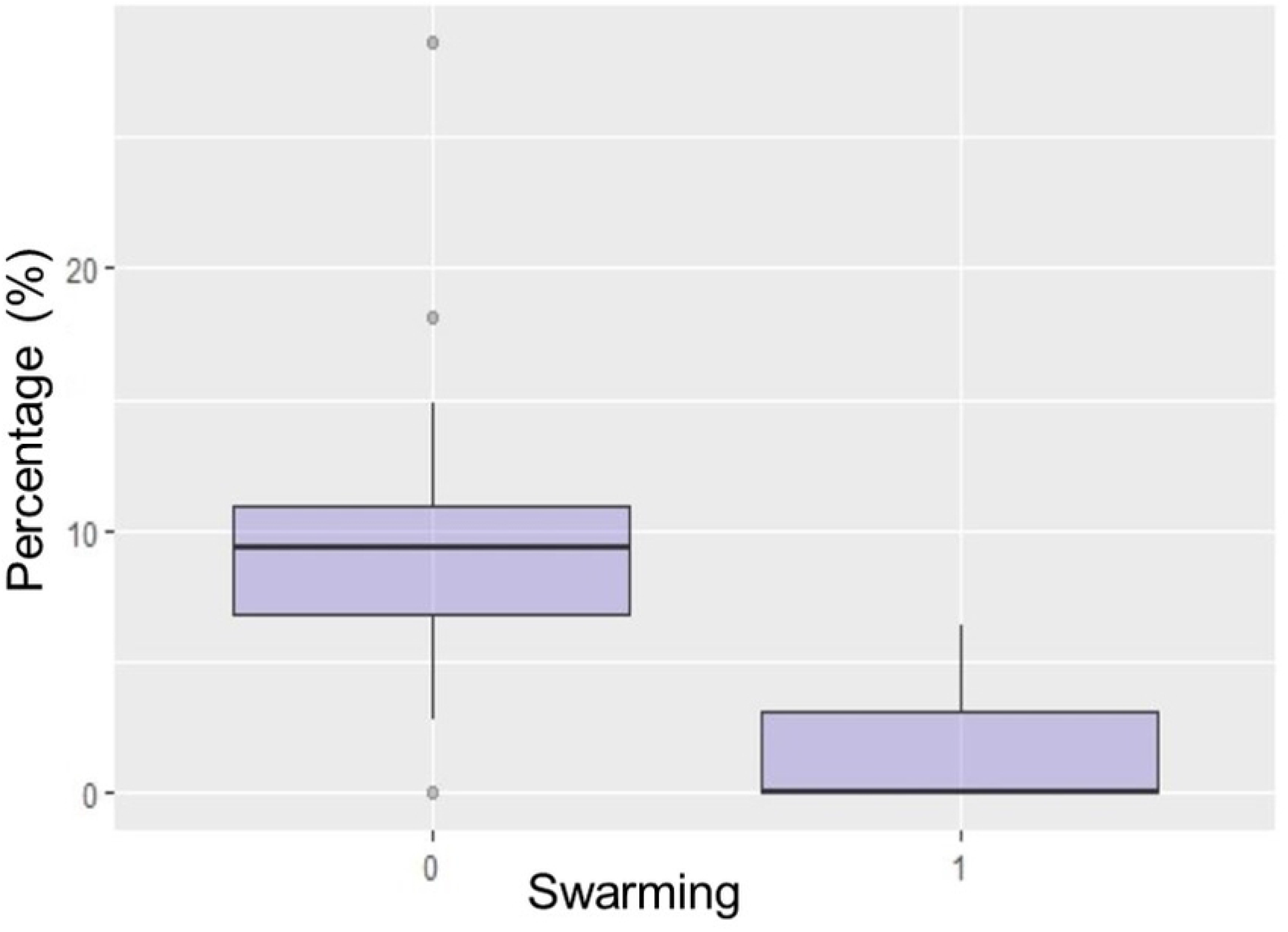
Study sites in Spain where bats were screened for AdVs. Blue dots indicate localities with at least one AdVs positive samples, orange dots indicate localities with negative samples for AdVs. (Modified from HansenBCN Miguillen - Own work, Public Domain, https://commons.wikimedia.org/w/index.php?curid=10577766)

Data from Vidovszky *et al.* [26] and Sonntag *et al.* [25] included. Lower and upper limits of 95% confidence intervals of the percentage have been included.

The number of individuals sampled within species varied considerably and 5 species (*Myotis daubentonii, Miniopterus schreibersii, Nyctalus lasiopterus, Pipistrellus kuhlii, P. pygmaeus*), accounted for 50% of the total number of individuals sampled for this study. On the other hand, the species *Eptesicus nilssonii, Myotis alcathoe, M. brandtii, M. crypticus, Rhinolophus mehelyii* were represented by fewer than 10 individuals and were consequently discarded from the proportion-based analyses of prevalence. Values of proportion of positive samples varied considerably across species, averaging 7.02% (Table 1). The species targeted for the individual-level analyses were: *Nyctalus lasiopterus* (10.26% prevalence), *Pipistrellus pygmaeus* (10.07% prevalence) *and Pipistrellus kuhlii* (9.57% prevalence).

**Table 1:**
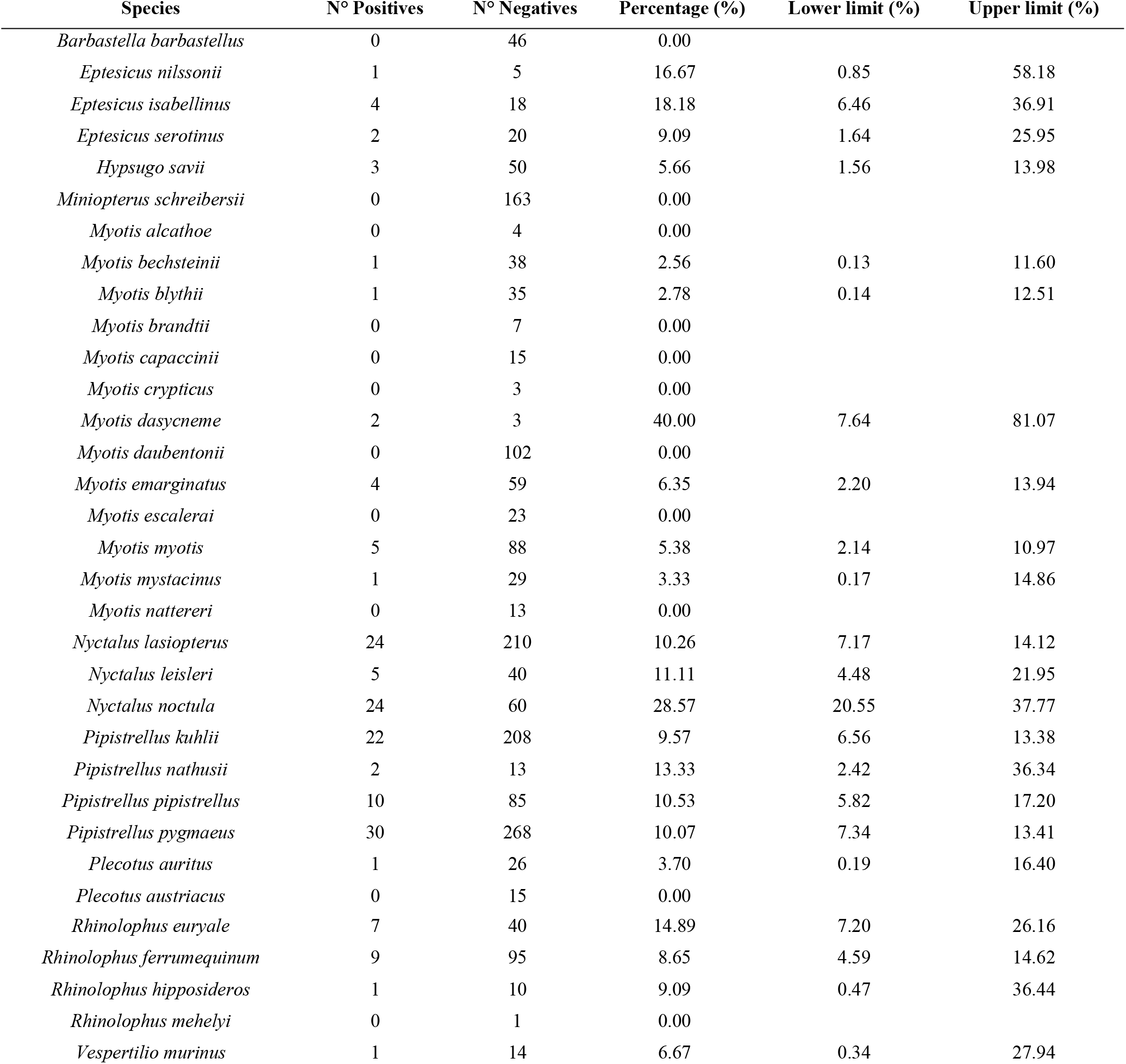
Number of individuals screened as positive or negative for Adenoviruses and percentage of positives for each species taken into account in this study.

### Phylogenetic Signal

The fully resolved phylogeny for all the studied species with proportional branch lengths was used to estimate the different phylogenetic component in AdVs presence (S1 Fig). Fritz’s *D* parameter was 0.482, and we detected a marginally significant departure from Brownian motion structure (*p* =0.058), whereas the probability of *D* resulting from Brownian motion structure was *p* = 0.165. The value of Pagel’s *λ* when analyzing the distribution of proportion of AdVs across the phylogeny was *λ* = 0.99 and highly significant (*p* =0.0003) and Blomberg’ *K* was *K* = 0.74 (*p* =0.002). Thus, the distribution of the AdVs infection across the studied European bats showed a strong phylogenetic component following Brownian motion.

### Trait association with AdVs prevalence

The phylogenetic linear regression analysis showed that regardless of the phylogenetic model used (although Brownian motion had the lowest AIC score, all three models tested were within ΔAIC ≤ 3), the variable MATING STRATEGY showed a significant (and negative) effect, as shown in Fig 2 (model=Brownian motion, *t* = −2.8920, *p* = 0.0093; all model results presented in S1 Table). Moreover, the same result was recovered when using AdVs presence as a binomial dependent variable (S2 Table), indicating a significantly higher AdVs presence in bat species not engaging in swarming for mating.

**Fig 2.**
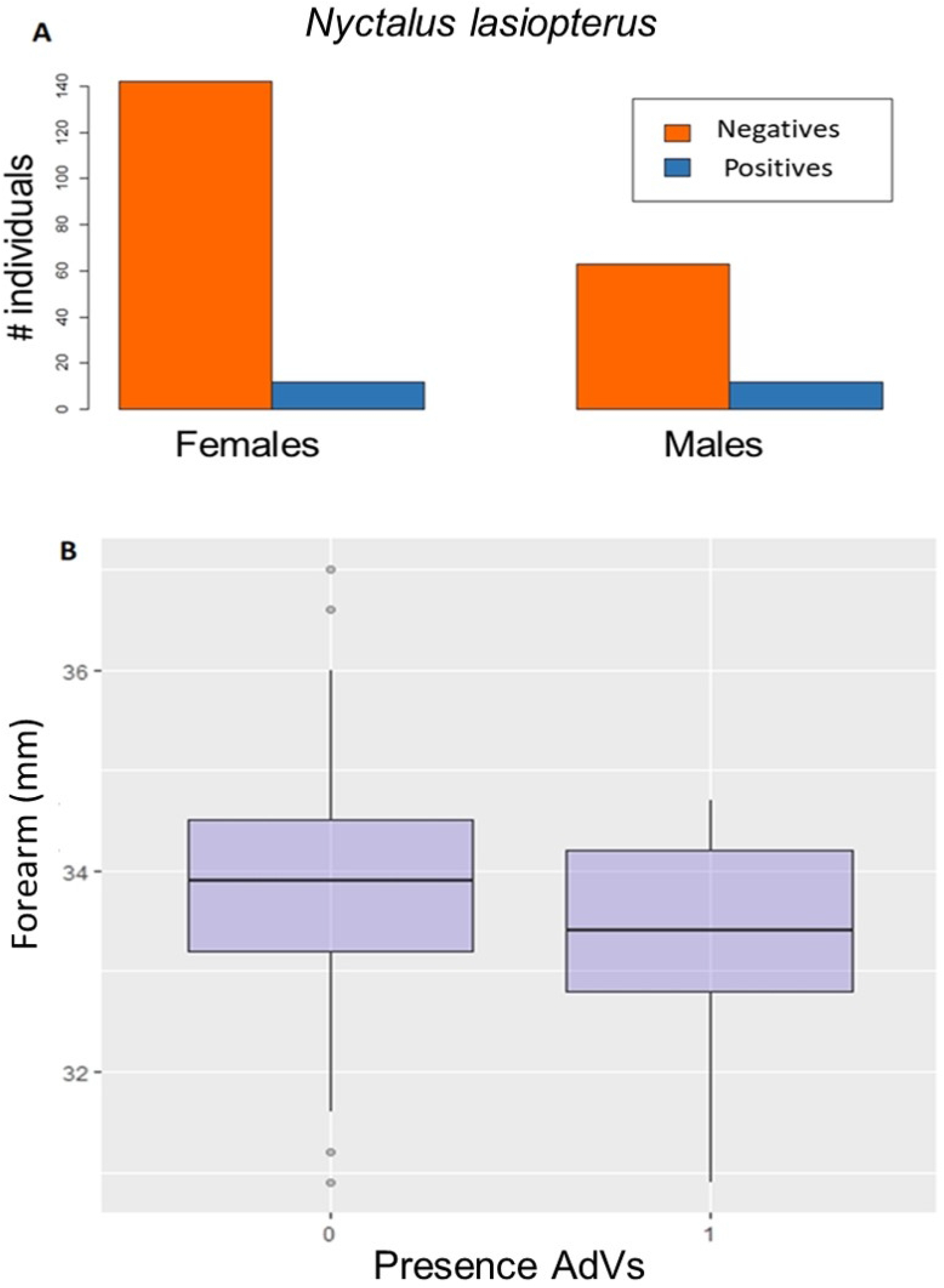
Boxplot of the percentage of positive individuals for species engaging in swarming and species not engaging in swarming. 0 indicates species not using swarming as mating strategy, 1 indicates species using swarming as mating strategy. (*N*=160).

At individual level and for the three species considered, the complete models were used since no model was selected by AIC (ΔAIC < 3). The GLMM analysis revealed a significantly higher AdVs presence for males (*z*= 2.067, *p*=0.039) in *N. lasiopterus* (Fig 3A), and a trend towards higher AdVs prevalence for individuals with smaller forearm (Fig 3B) for *P. kuhlii* (*z*= - 1.656, *p*= 0.098). The variance of the model explained by only the random variable was 4% for *N. lasiopterus*, 62% for *P. pygmaeus* and 16% for *P. kuhlii*. Results for all the variables selected are available in supplementary material S3 Table.

**Fig 3.**
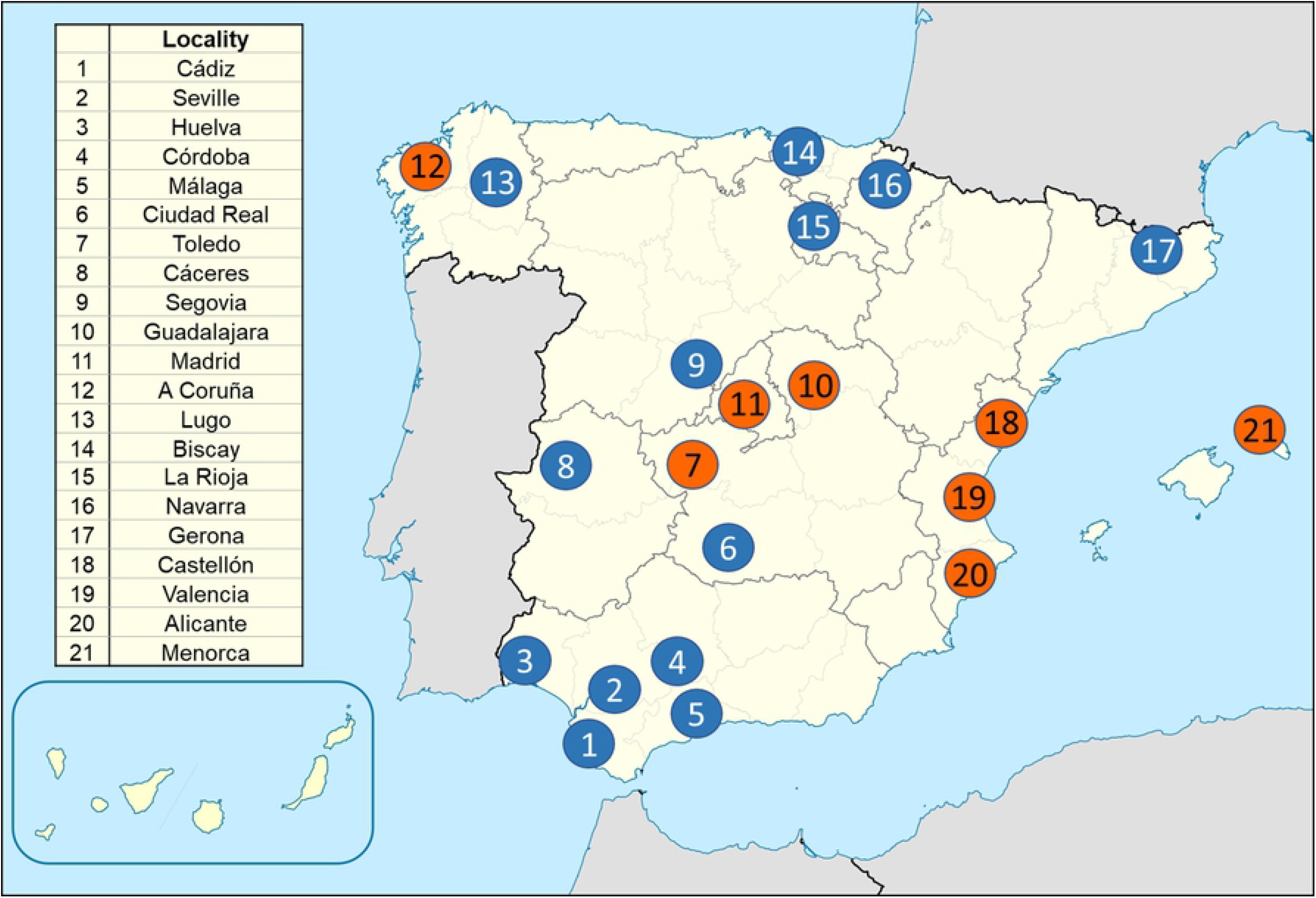
Results of within-species analysis. (**A**) Barplot by sex of number of individuals positive (blue) and negative (red) to Adenovirus, for *Nyctalus lasiopterus*. (**B**) Boxplot for individuals screening negative for AdVs (0) and positive for AdVs (1) in relation to forearm length for *Pipistrellus kuhlii*.

### Mantel test

The Mantel tests showed no significant correlation for all pairwise comparisons between geographic distances and differences in the proportion of AdVs presence across localities for *N. lasiopterus* (*r* = −0.3186; *p* = 0.941) and *P. kuhlii* (*r* =0.1132; *p*= 0.113) (S2 Fig), but this correlation was significant for *P. pygmaeus* (*r* = 0.3364; *p*= 0.036) indicating that with greater distance, there will be a greater difference in the proportion of individuals affected by AdVs (S2 Fig).

## Discussion

For the first time to our knowledge, Adenovirus prevalence and its correlation with host traits is studied across a wide range of bats both at the species and individual level. The understanding of which and how host traits affect the presence of viruses in bats is a key step to the understanding of the transmission mechanisms and evolutionary strategies of viruses. Such mechanisms are key in the process of host switching that cause the appearance of emerging diseases and therefore its understanding is important to improve disease managements [35]. Our results show that Adenoviruses seem to be quite common in European bats since they were found across all the studied tribes and in most genera (except for *Barbastella* and *Miniopterus*) but their prevalence varied considerably among species [16]. The three most abundant species showing the highest frequency of adenovirus infection (*Nyctalus lasiopterus*, *Pipistrellus pygmaeus* and *P. kuhlii*) experienced an average of 9.97% prevalence. In contrast, other species showed very low Adenovirus presence and was completely absent in others despite their large sample sizes (e.g.: *Miniopterus schreibersii*) but in other species, the absence of positive results could be due to the low number of sampled individuals (generally the rarest bats in Europe).

Fritz’s D, Pagel’s *λ* and Blomberg’s *K* parameters all pointed to a strong phylogenetic signal in the presence of AdVs across European bats and consequently, the need for accounting for phylogenetic relationships in all subsequent models. This phylogenetic component of the distribution pattern of AdVs in bats was also recently suggested by Iglesias-Caballero *et al.* [16] as the sequences of new mastadenoviruses were clustering generally in agreement with the host bat families or even with the bat species.

Contrary to a previous study on the factors influencing viruses on bats [32], we have not found significant effect of the bats’ group size on the presence of Adenoviruses for European species. In their study, Webber *et al.* [32] had a wider perspective and focused on overall viral richness in bats, whereas this study is centered exclusively on AdVs and so our differing results may indicate that Adenoviruses use different transmission pathways than other viruses. Transmission of respiratory AdVs in humans requires close contact although it can possibly occur through droplet spray (such as those produced by coughing or sneezing) and/or aerosols, but data are still limited [27–29]. In general, little is known about transmission mechanisms of Adenoviruses and to our knowledge, no study has focused on this particular aspect across bats, although the host species specificity found for most of the Adenoviruses in bats [16] points to cross-species contacts as rare events.

Counterintuitively, species engaging in swarming behavior were found to have significantly lower prevalence of Adenoviruses than other bats despite having theoretically greater chances of contact. Swarming related to mating is shown mainly by forest species and is described as the gathering of bats, commonly in caves and underground sites, during a few hours after dusk and for a few days [36]. Two hypotheses could explain this finding. Firstly, bat species engaging in swarming seem to be in contact for only a short time, which would imply a lower chance of transmission compared to bats with polygynous mating systems, like harems, which stay in contact for longer periods of time, sometimes all year around [37]. A second plausible hypothesis is based on the fact that mammals with high infection risk, especially those with promiscuous behavior, develop a stronger immune system compared to species with low infection risk [38,39]. Thus, it seems reasonable that bats with swarming behavior may show stronger immune response protecting them from infections, despite being very costly and even represent a trade-off with other life-history traits [39].

When analyzing the effect on the AdVs presence of within-species variance in traits, we found that the site of capture explains an important part of the variance of all models suggesting an underlying general pattern across species and supporting the contact-rate hypothesis. This trend is particularly clear for *Pipistrellus pygmaeus*, whose sampling localities are more widespread along the Iberian Peninsula than those for *Nyctalus lasiopterus* and *P. kuhlii* (Fig 4).

**Fig 4.**
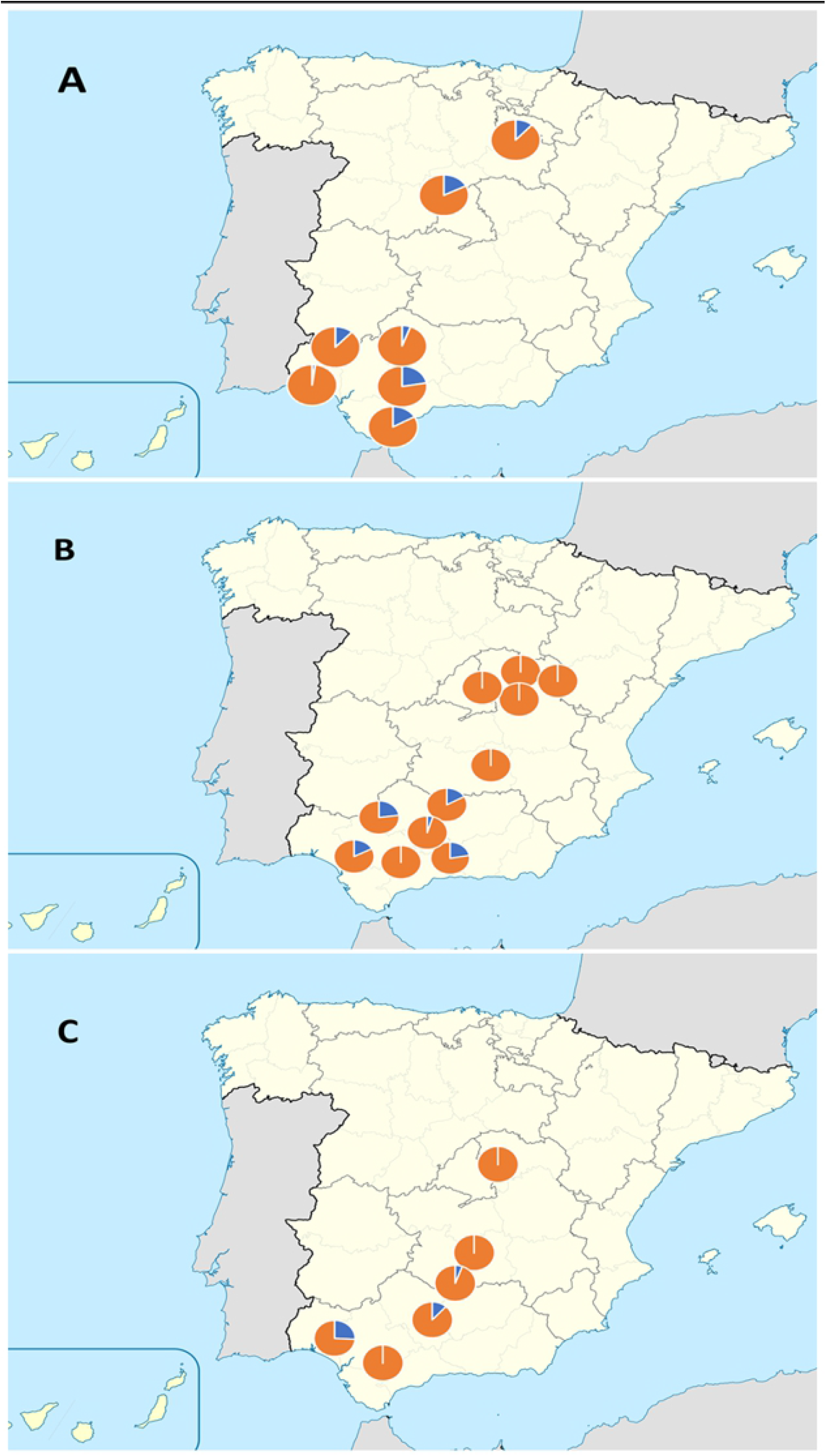
Sampling localities with percentage of AdVs positive samples. Portion of AdVs positive sample is indicated in blue, AdVs negative samples in orange for (**A**) *Nyctalus lasiopterus*, (**B**) *Pipistrellus pygmaeus* (**C**) *Pipistrellus kuhlii*. Only localities with at least 5 recorded individuals are shown. (Modified from HansenBCN Miguillen - Own work, Public Domain, https://commons.wikimedia.org/w/index.php?curid=10577766)

Following the contact-rate hypothesis, bats from a specific locality interact with other individuals of the same area, favoring the virus transmission locally [7]. A similar pattern was already found for the viruses responsible for avian influence, Marburg or Ebola [40–42]. In this direction, the Mantel’s test supported a significant geographic component in the incidence of the Adenoviruses for *Pipistrellus pygmaeus* which is typically sedentary species [34].

A higher prevalence of Adenoviruses was found in males for *Nyctalus lasiopterus*, the species in which the different sexes were best represented. This result is particularly interesting since it could affect the general dynamics of the virus given the generally predominant role played by males in dispersal [43]. The possibility of a sex byes infection as a general pattern should be explored further in other bat species and could be related to the trade-off between immune function and reproduction, making males more susceptible to virus infections during reproductive season due to the immune-suppressant effect of testosterone production [44].

Our results show that, individuals with smaller forearm are more likely to be infected by Adenoviruses for *Pipistrellus kuhlii*. Subadult bats usually have slightly longer forearms [34] and the lower prevalence in bats with larger measures could be explained by the influence of the antibody loads transferred to them through their mother’s milk in still developing bats, making them less suitable to Adenoviruses prevalence [45]. This is just a possibility and further studies should help understand the relationships between Adenoviruses presence and age.

In sum, our findings highlight a common occurrence of Adenoviruses in European bats and provide evidence of the importance of mating behavior in the prevalence of Adenoviruses, contrary to what has been suggested generally for viruses. Besides, the three European bats species that had higher Adenovirus prevalence do not show any common pattern, pointing to Adenoviruses transmission as a complex process. Our study highlights the importance of combining behavioral with ecological traits in explaining viral richness and transmission.

## Materials and Methods

### Ethics Statement

Non-lethal sampling was based on permits 201710730002961/IRM/MDCG/mes issued by Dirección General de Gestión del Medio Natural y Espacios Protegidos (Consejería de Medio Ambiente, Junta de Andalucía, Spain), 10/085545.9/17.9/17 issued by Consejería de Medio Ambiente, Administración Local y Ordenación del Territorio, Comunidad de Madrid, and PNSNG_SG_2018_0093 issued by Servicio Territorial de Medio Ambiente de Segovia, Junta de Castilla y León. The techniques used meet the guidelines published by the American Society of Mammalogists on the use of wild mammals in research [46]

### Data collection

Most of the information used in this study was obtained during a surveillance program for bat rhabdovirus and lyssavirus carried out between 2004 and 2008, 2016 and 2018. Bats were mist-netted near roosts or over water and released in the same collecting point after sampling. Each animal was identified, sexed, measured and weighed. Sampling consisted of obtaining a membrane wing-punch, saliva with oropharyngeal swabs and stool samples (when this was possible). Swabs were stored in 1.5 mL tubes filled with lysis buffer. All samples were aliquoted and stored at −80 °C prior to analysis. Samples were screened for AdVs following published protocols [16,47,48]. Our own database was completed with the published data of AdVs presence in bats from Germany and Hungary [25,26]. The identification of bats belonging to species complexes was confirmed through PCR amplification of a diagnostic mtDNA fragment following Ibañez *et al.*, [49] and Kaňuch *et al.* [50].

### Phylogeny and phylogenetic signal

A fully resolved phylogeny containing all the study species was not available from the literature and is notoriously difficult to reconstruct for this group, [51] and so a hybrid approach was taken whereby, published trees were grafted to obtain a full-solved topology whereas branch lengths were estimated using molecular sequence data. The published phylogenetic studies for the four families existing in Europe, used to manually construct a topology of all species in this study were Guillén-Servent *et al.*, [52] for relationship in the family Rhinolophidae; Ruedi *et al.*, [53] and Stadelmann *et al.* [54] for the subfamily Myotinae within Vespertilionidae; and Hoofer and Bussche [55] for the rest of this last family and for the family Miniopteridae. Complete sequences of the mitochondrial genes *cytochrome b (CYTB)* and *NADH dehydrogenase 1 (ND1)*were obtained from GenBank for all available European species and were used to estimate branch length in the constructed topology. This dataset needed to generate sequences *de novo* for the species *Miniopterus schreibersii* for which these markers were not available. The new sequences are deposited in GenBank under the accession numbers MK737740 and MK737741. For each locus sequences were aligned with the software ‘ALTER’ from http://darwin.uvigo.es/software/ [56] and MEGA [57], and partitionfinder v2.1.1 [58] was run on the concatenated alignment (1805 base pairs in length) to obtain the optimal partitioning scheme and substitution models. Partitionfinder was set to use unlinked branch lengths, search only BEAST models with the ‘greedy’ algorithm and AICc as the model selection criterion. A maximum of six possible partitions were allowed; the three codon positions of the two loci. BEAST v2.4.7 [59] was then fed the alignment and the fixed topology, and using a relaxed clock log normal model with exponential priors on the mean (mean=10) and standard deviation (mean=0.33), was allowed to estimate the branch lengths over 20 million MCMC iterations, storing every 1000th. Chain diagnostics was performed using Tracer v1.7 [60] to ensure sufficient mixing and parameter convergence and a maximum clade credibility tree was generated using TreeAnnotator v.2.4.7 (part of BEAST package) using a 10% burn-in and median node heights.

Fritz’s *D*, Pagel’s *λ* and Blomberg’s *K* parameters were used to estimate the strength of the phylogenetic signal affecting the pattern of AdVs presence across European bats. Fritz’s *D* was used for binary data (presence/absence) so that values close to one indicate that the distribution of the binary trait is random with respect to the given phylogeny and values close to zero indicate the trait is distributed as expected under a Brownian motion model of evolution [61]. When the variable estimated was the proportion of infected bats in each species, Pagel’s *λ* and Blomberg’s *K* parameters were used. Values of *λ* range from zero to one where *λ* = 0 indicates that related species do not share similar values for the trait (percentage of infection) and *λ* = 1 indicates a pattern fully explained by the phylogenetic relationships under Brownian motion, with related species showing similar values for the given trait. Finally, *K* is scaled so that zero indicates no phylogenetic signal, K=1 is the expected value for trait evolution under Brownian motion and values higher than one suggest stronger phylogenetic signal than predicted by Brownian motion [62,63].

### Variables Selection and modelling

Ecological and behavioural traits were selected considering their *‘a priori’* importance for the virus transmission and their availability for the European bat species from the bat literature and general revisions [64], identification guides [34] (S4 Table). For the comparative analysis across all European bats, the following variables were recorded for each species: (1) GROUP SIZE, defined as the upper bound of individuals found usually in summer roosts; (2)

FOREARM: defined as the average range of forearm length (in mm) reported for each species; (3) SOCIABILITY defined as the chance of sharing summer roost with other species and considering just two categories: species that have never been found sharing their roost with other species and species that sometimes or always share roost with other species; (4) MATING STRATEGY: considered as whether mating takes place in seasonal swarming or not; (5) MIGRATION: defined as whether the bat species is known to perform long-distance movements (longer than 100 km), regional (between 10 and 100 km) or no movements; (6) ROOST TYPE: considered in three categories: roosts in caves, trees or crevices. At the individual level, the analyses focused on the three species that showed the highest AdVs presence combined with a large enough sample size (> 200 individuals). The recorded variables were: (1) LOCALITY: site of capture; (2) SEX: male or female; (3) FOREARM: Measured in mm.

For among-species comparisons, AdVs prevalence was analyzed as the arcsin-transformed proportion of positive samples over the total number of samples for each species. Species with fewer than 10 samples were eliminated from the matrix and from the phylogeny. The analyses were also run considering presence/absence of AdVs as a dependent variable and including in this case all sampled species. The analyses were performed with R statistical computing packages [65] and scripts are available upon request. Fritz’s D, Pagel’s λ and Blomberg’s K parameters were estimated with the packages ‘*Caper*’ [66] and ‘Phytools’[67]. Phylogenetic linear regressions were carried out with the package ‘*Phylolm*’ [68] that respects the shared evolutionary histories of species [69]. Models of trait evolution were compared assuming correlation structures under either Brownian motion, Ornstein-Uhlenbeck or Pagel models [70] and the best model was selected based on the Akaike Information Criterion (AIC) [71]. As a threshold value for model selection, a model showing ΔAIC > 3 was taken as having greater support [72]. Complementary, a binary phylogenetic generalized linear model was run for each species, with presence of AdVs as the dependent variable and using the ‘*Ape*’ package [73].

Within species, we used general linear mixed models implemented in the ‘*Lme4*’ package [74] to test for associations between traits and presence of AdVs, and considering AdVs presence as a binary dependent variable. In order to account for uncontrolled spatial variation, LOCALITY was included in the models as a random variable. The best linear mixed model was again selected based on AIC and same threshold criterion. As a confirmatory analysis for the models, the variance explained by only the fixed variables was compared to the variance obtained including the random variables using the package *‘Mumin’* [75]

At both among-species and within-species levels, inter-correlation was were checked using a Pearson’s correlation test for continuous variables and an ANOVA for categorical variables. The spatial distribution of the AdVs positive bats was inspected by performing a Mantel test between the matrix of geographic distances and the matrix of differences in proportion of infection between sites with the package *‘Ade4’* [76,77].

## Acknowledgments

We are grateful to the bats team of the Estación Biológica de Doñana (CSIC) and to Francisco Pozo and collaborators of the Instituto de Salud Pública Carlos III for their collaboration. We thank Jens Rydell, David Serrano, Josué Martinez de la Puente, Josean Donazar and Carlos Herrera for their useful suggestions and comments.

## Supporting information

**S1 Table. Result of the percentage models**. Models under Brownian motion, Pagel and Ornstein-Uhlenbeck structures.

**S2 Table. Results of absence/presence model**. Estimate value, standard error, Z-score and p-value included.

**S3 Table. Result of GLMM model (individual level) for each species taken into account**. A: *Nyctalus lasiopterus*. B: *Pipistrellus pygmaeus*. C: *Pipistrellus kuhlii*.

**S4 Table. Traits used in the model for each species**. Information taken from: 1: Dietz *et al.* [34], 2: Action Plan for the Conservation of All Bat Species in the European Union [64]. Values indicated with asterisk have been obtained by personal communication of Jens Rydell and Jesús Nogueras Montiel.

**S1 Fig. Evolutionary relationships of bat species considered in this study**. Based on the phylogenetic hypotheses of: Guillén-Servent *et al* [52].; Hoofer and Bussche [55]; Ruedi *et al.* [53] and Stadelmann *et al.* [54]. Branch lengths reflect mitochondrial sequence divergences. Tip labels indicate species screening positive (black) or negative (white) for AdVs.

**S2 Figure. Scatterplot showing the results of the Mantel test**. Mantel test between the matrix of differences in percentages of AdVs presence and the matrix of geographic distances for (A) *Nyctalus lasiopterus* (B) *Pipistrellus pygmaeus* and (C) *P. kuhlii*.

